# An allele-specific functional SNP associated with two autoimmune diseases modulates *IRF5* expression by long-range chromatin loop formation

**DOI:** 10.1101/533661

**Authors:** Hlaing Nwe Thynn, Xiao-Feng Chen, Wei-Xin Hu, Yuan-Yuan Duan, Dong-Li Zhu, Hao Chen, Nai-Ning Wang, Huan-Huan Chen, Yu Rong, Bing-Jie Lu, Man Yang, Feng Jiang, Shan-Shan Dong, Yan Guo, Tie-Lin Yang

## Abstract

Both Systemic Lupus Erythematosus (SLE) and Systemic Sclerosis (SSc) are autoimmune diseases sharing similar genetic backgrounds. Genome-wide association studies (GWASs) have constantly disclosed numerous genetic variants conferring to both disease risks at 7q32.1, but the functional mechanisms underlying them are still largely unknown. Through combining fine-mapping and functional epigenomic analyses, we prioritized a potential independent functional SNP (rs13239597) within *TNPO3* promoter region, residing in a putative enhancer element. Functional analysis integrating expression quantitative trait locus (eQTL) and high-throughput chromatin interaction (Hi-C) demonstrated that *IRF5* is the distal target gene (~118kb) of rs13239597, which is a key regulator of pathogenic autoantibody dysregulation increased risk of both SLE and SSc. We experimentally validated the long-range chromatin interactions between rs13239597 and *IRF5* using chromosome conformation capture (3C) assay. We further demonstrated that rs13239597-A acted as an allele-specific enhancer regulating *IRF5* expression, independently of *TNPO3* by using dual-luciferase reporter assays and CRISPR-Cas9. Particularly, the transcription factor EVI1 could preferentially bind to rs13239597-A allele and increase the enhancer activity to regulate *IRF5* expression. Taken together, our results uncovered the mechanistic insight connecting between a noncoding functional variant with a distal immunologically pathogenic gene *IRF5*, which might obligate in understanding the complex genetic architectures of SLE and SSc pathogenesis.

## Introduction

Systemic Lupus Erythematosus (SLE [MIM: 152700]) and Systemic Sclerosis (SSc [MIM: 181750]) are two typical systemic autoimmune diseases, which share pathogenic features such as interferon signature, loss of tolerance against self-nuclear antigens, multi-tissue damage and platelet system activation [1]. The worldwide prevalence is about 32 for SLE and 24 for SSc per 100,000 [2]. Although the etiology of SLE and SSc remains unclear, genetic factors are considered as the key query point. Genome-wide association studies (GWASs) have identified a large number of risk genes associated with SLE and SSc, some of which are pleiotropic genes for both diseases such as *STAT4* (MIM: 600558) [3, 4], indicating that these two diseases share a portion of the genetic backgrounds.

Chromosome 7q32.1 locus harboring *IRF5-TNPO3* (*IRF5* [MIM: 607218], *TNPO3* [MIM: 610032]) has been reported repeatedly as a strong association signal with SSc or SLE outside the human leukocyte antigen (*HLA*) region [5, 6]. This *IRF5-TNPO3* region also contains some susceptibility variants associated with other auto immune disorders, like rheumatoid arthritis, Sjogren’s syndrome and multiple sclerosis [7]. A pan-meta GWAS study combing both SLE and SSc samples found the highest association signal for the SNP rs13239597 (*P* = 1.17 × 10^−29^) [8]. This SNP is located in the promoter region of *TNPO3* (Transportin 3) and ~118 kb downstream of *IRF5* (Interferon regulatory factor 5). *IRF5* is a well-known immunologic gene with crucial regulatory roles in modulating immune responses across numerous immune-related cell types in Toll-like receptor (TLR) signaling pathway [9]. However, the functional roles of *TNPO3* in autoimmune etiology are largely unknown. Recent studies have found that SNPs within the potential regulatory elements could regulate expression of distal genes by long-range interactions [10, 11]. It is important and interesting to decipher the functional roles of these autoimmune disease-associated SNPs and find out whether *TNPO3* or *IRF5* is the true target gene regulated by these variants, which might help fulfill the gap between GWAS findings and autoimmune disease etiology.

Therefore, in this study, we firstly prioritized a potential functional independent variant (rs13239597) located in an enhancer element at 7q32.1 through combining genomic fine-mapping and functional epigenomic analyses. Then, we optimized *IRF5* as the distal target gene for rs13239597 by integrating expression quantitative trait locus (eQTL) and high-throughput chromatin interaction (Hi-C) analysis. Such long-range chromatin interactions was also validated by using chromosome conformation capture (3C) assay. We further demonstrated that rs13239597 could act as an allele-specific strong enhancer to regulate *IRF5* expression through a series of functional assays including dual-luciferase reporter assays and CRISPR-Cas9. Finally, we found that transcription factor EVI1 (MIM: 165215) could preferentially bind to rs13239597-A allele to increase the enhancer activity on *IRF5* expression. Our findings uncovered a novel functional mechanisms connecting SNPs at *TNPO3* locus with *IRF5* in a long-range chromatin regulatory manner, which would provide promising routes towards the improved multidisciplinary therapy of complex autoimmune diseases.

## Materials and Methods

### Conditional association and Bayesian fine-mapping analysis

Genome-wide association analysis results on SLE at 7q32.1 locus were downloaded from a recent large-scale meta-analysis comprised 23,210 European samples [12]. Linkage disequilibrium (LD) analyses were conducted using Haploview Version 4.2 [13] in European samples from the 1000 genome V3 genotype data [14]. To identify potential independent SLE-associated SNP(s), we performed a stepwise conditional association analysis using GCTA-COJO (--cojo-slct) [15, 16] analysis with LD correlations between SNPs estimated from 8748 unrelated samples from the Atherosclerosis Risk in Communities (ARIC) data (dbGap: phs000280.v3.p1.c1) [17]. To clarify the independent SLE association signal surrounding *TNPO3* locus, we further performed association analysis of *TNPO3* locus SNPs by conditioning on identified independent SNPs in *IRF5* locus. To prioritize potential causal SNPs surrounding *TNPO3* locus, we performed Bayesian fine-mapping analysis to identify 95% credible SNP set, which was originally described by Maller *et al*. [18]

### Functional epigenetic annotation

To prioritize potential functional variants, we annotated several enhancer related epigenetic markers for the regions surrounding our interest SNPs using ChIP-seq data from ENCODE [19], including the DNase I hypersensitivity (DHS), the histone markers (H3K27ac, H3K4me3), RNA polymerase II (Pol 2) and histone acetyltransferase p300 binding sites in immune related blood cell lines such as GM12878 lymphoblastoid cell line, primary T cells and primary B cells respectively. All annotated data were visualized by using WashU Epigenome Browser (v46.1).

### Cis-eQTL analysis

Matched SNP genotype and RNA-seq data for 373 unrelated human lymphoblastoid cell lines (LCLs) samples of European population were retrieved from 1000 genome V3 genotype data [14] and ArrayExpress (E-GEUV-1) [20] respectively. We transformed genotype data into plink format using vcftools. By using the linear regression model implemented in PLINK, cis-eQTL analysis was conducted between selected SNPs and expression of nearby genes within 1 Mb region. For further validation, cis-eQTL association from publicly available Genotype-Tissue Expression (GTEx) database, encompassing over 25,000 samples from 714 donors across 53 tissues was also checked [21] with corresponding genotype data obtained from dbGaP (phs000424.v7.p2). We next queried for cis-eQTL in another peripheral blood samples from 5,311 individuals [22].

### Hi-C and TAD analysis

To validate the long-range regulation between focused SNP and its eQTL target gene, we checked their chromatin interactions using Hi-C or capture Hi-C data on GM12878 and CD34 cells downloaded from the previous studies [23, 24]. Hi-C data in IMR90 cells were obtained from 4D Genome databases [25, 26]. The original ChIA-PET data and newly improved ChIA-PET data on six cell lines (K562, NB4, HCT-116, Hela-S3, GM12878, and MCF7) were acquired from the UCSC ENCODE download portal [27]. BEDTools [28] was used to extract the chromatic interacted regions. We further checked whether the focused SNP and its target gene are within the same topologically associating domain (TAD) region using TAD data in IMR90 cells collected from GEO database (GSE35156) [29].

### Motif analysis

We conducted motif analysis surrounding rs13239597 (50-bp) using MEME Suite toolkit [30] with three publicly available TF motif databases, namely JASPAR [31], HOCOMOCO [32], and SwissRegulon [33]. The motif with allele-specific binding at rs13239597 was retained.

### Comparison of *IRF5* expression between SLE patients and healthy samples

We retrieved three SLE genome-wide gene expression datasets in whole blood samples from the Gene Expression Omnibus (GEO) repository, including 30 healthy controls and 99 SLE patients for GSE61635, 46 healthy controls and 96 patients for GSE39088 and 72 healthy controls and 924 patients for GSE65391[34]. We compared the average expression level for all microarray probes on *IRF5* between SLE patients and healthy samples in each dataset using two-tailed student’s T test, respectively.

### Culture of cell lines

The EBV-transformed B lymphocyte cells (Raji), the human embryonic kidney 293T cells (HEK293T) and the human bone osteosarcoma epithelial cells (U2OS) were purchased from the American Type Culture Collection (ATCC, USA) and cultured in RPMI-1640 (Roswell Park Memorial Institute Medium) medium for Raji and U2OS cells and DMEM (Dulbecco’s Modified Eagle’s Medium) medium for HEK293T cells supplemented with 10% fetal bovine serum (FBS) (Invitrogen, USA), 100U/mL penicillin and 0.1 mg/mL streptomycin in 5% CO_2_ at 37°C incubator.

### Genotyping of rs13239597

To measure genotypes of rs13239597 over Raji, HEK293T and U2OS cell lines, we extracted the human genomic DNAs from each cell line and amplified the fragment surrounding rs13239597 using the same primers used in the Luciferase expression plasmid constructs (Table S3). The amplified DNAs were digested with the restriction enzyme EcoRV, followed by the 1% gel electrophoresis analysis and further confirmed by sequencing.

### Luciferase expression plasmid constructs

A 1000-bp putative enhancer fragment containing the reference or alternate allele of rs13239597, a 1323-bp promoter fragment surrounding *IRF5* transcription start site, as well as a 1137-bp promoter fragment surrounding *TNPO3* transcription start site were amplified using PCR from healthy human genomic DNA (Table S3). The amplified enhancer and *IRF5* promoter fragments were ligated and cloned into the enzyme cut sites MluI and HindIII of pGL3-basic vector. The individual *IRF5* or *TNPO3* promoter fragment was cloned into the enzyme cut sites SmaI and HindIII or XhoI and HindIII of pGL3-basic vector as the baseline control, respectively. We also amplified a 1698-bp fragment including both rs13239597 enhancer and *TNPO3* promoter and cloned it into the enzyme cut sites MluI and HindIII of pGL3-basic vector (Table S3). The site-directed mutagenesis was used to generate the constructs containing the other allele not amplified from initial cloning with the Quick Change II Site-Directed Mutagenesis Kit (Agilent Technology, USA). All the constructed plasmids were validated by sequencing and did not contain any other sequence variations. The primers used in these constructs were listed in Table S3.

### Transfection, electroporation and dual-luciferase reporter assays

The constructed expression plasmids were transfected into HEK293T and U2OS cells by using ViaFect^™^ transfection reagent (Promega, USA) according to the manufacturer’s instructions. Celetrix electroporation method was used to transfect the expression plasmids into Raji cells according to the manufacturer’s instructions using the electroporation machine (Catalog# CTX-1500A LE), the pressured electroporation tubes (Catalog# 12-0107), and the electroporation buffer (Catalog# 13-0104) (Celetrix LLC, USA). 5-8 million of Raji cells were used to transfect 2 μg of expression plasmids by using 320 volt. An internal control reporter vector, pRL-TK containing *Renilla* luciferase (Promega, USA), was simultaneously transfected into the cells with expression plasmids as an internal control for assay-to-assay variability. And then, the transfected cells were incubated in 5% CO_2_ at 37°C incubator. After 48h of transfection, the cells were harvested and investigated for luciferase activity using the Dual-Luciferase Reporter Assay System (Promega, USA). Luciferase activity was normalized through division of major or minor allele construct luciferase signals by pRL-TK luciferase signals. The mean and standard error of measurement were calculated on the basis of the normalized luciferase activities. The results were obtained from three independent experiments and each experiment was done in triplicate.

### shRNA expression plasmid constructs and shRNA knockdown

For shRNA knockdown of transcription factor EVI1 and *TNPO3*, the two independent miR-30-styled short hairpin RNAs (shRNA-1 and shRNA-2) expression plasmids were constructed by using EVI1- or *TNPO3*-targeted sense and antisense oligonucleotides. Each pair of sense and antisense oligonucleotides were connected with miR30 backbone according to the previous protocol [35]. Each resulted miR-30 styled shRNA was amplified with the primers shown in Table S3 and purified to be cloned into XhoI and EcoRI enzyme sites of pcDNA3.1 vector (Invitrogen, Carlsbad, USA). For the negative control (NC), shRNA NC expression plasmid was also constructed in the same way. 2 μg of each plasmid (shRNA 1, shRNA 2 and shRNA NC) were independently transfected into 70-80% confluence of HEK293T cells or U2OS cells by using ViaFect^™^ transfection reagent (Promega, USA) according to the manufacturer’s instruction. After 48h of transfection, total RNA was isolated to detect the mRNA expression by RT-qPCR. Moreover, EVI1 shRNA plasmids were independently co-transfected with the expression plasmid including rs13239597-C allele or rs13239597-A allele with *IRF5* promoter used in the luciferase reporter assay. The transfection and measuring of luciferase activity are the same as indicated in dual-luciferase reporter assay section. The results were obtained from three independent experiments and each experiment was done in triplicate.

### Enhancer deletion by CRISPR-Cas9

To efficiently eliminate the enhancer fragment (1000-bp) residing rs13239597, CRISPR-associated RNA-guided endonuclease Cas9 cleavage technology was used [36]. Briefly, we first designed a set of single-guided RNAs (sgRNAs) targeting 8 sites around upstream and downstream of the enhancer fragment by using the CRISPR design platform (V1 tool) maintained by the Zhang Lab at the Board Institute. Oligonucleotides containing these sgRNAs were cloned into the BsaI enzyme site of pUC19-hU6-sgRNA backbone vector and then, the same amount of each sgRNA plasmid with pCas9 plasmid (Addgene plasmid #42876) were co-transfected into HEK293T cells using ViaFect^™^ transfection reagent (Promega, USA) according to the manufacturer’s instruction. After 72h transfection, the genomic DNA were extracted, followed by PCR amplification of target fragment and T7 endonuclease I assay to quantify the indel (insertion and deletion) percentage as previously described. From these results, the best pair of sgRNAs including upstream sgRNA (491-bp far from rs13239597) and downstream sgRNA (444-bp far from rs13239597) was chosen and cloned together into KpnI and NheI enzyme cut sites of lentiCRISPR v2 plasmid (Addgene plasmid # 52961). This resulted plasmid transfection into HEK293T and U2OS cells and electroporation into Raji cells were performed the same as mentioned in Luciferase reporter assay. After selection with puromycin (2 mg/ml), the cells were harvested for DNA and total RNA extraction for further analysis. The results were obtained from three independent experiments and each experiment was done in triplicate. sgRNAs primers and PCR primers are listed in Table S3.

### Chromosome Conformation Capture (3C) Assay

The 3C assay was performed as described previously [37] in Raji and U2OS cell lines. Briefly, approximately 1×10^8^ cells were fixed with 1% formaldehyde for 10 min and stopped the fixation reaction by quenching with 2.5 M glycine. The cells were lysed using lysis buffer (10mM Tris-HCL-pH8.0, 10mM NaCL, 0.2% Igepal and autoclaved water) and 200 μl of 10 × proteinase inhibitors and fractionated using a Dounce homogenizer for nuclear fraction. After washing the pellet two times with 1ml of 1 × NEB buffer 2.1 (New England BioLabs (NEB), USA), the nuclear lysates were digested with 800U KpnI (NEB) overnight at 37°C and 950 rpm, followed by the inactivation with 86 μl of 10% SDS for 30 min at 65°C. The cross-linked digested DNA were re-ligated with each 800U T4 DNA ligase (NEB) in 7 ml of ligation cocktail including 1.1 × ligation buffer (NEB) and 10% Triton X-100 for 2 hours at 16°C. The ligated samples were treated two times with fresh 10mg/ml proteinase K solution per tube and incubated at 65°C for 4 hours and overnight respectively. DNA was extracted with phenol-chloroform and precipitated with ethanol-acetate. Quantification of 3C interaction products were undertaken by PCR followed by agarose gel electrophoresis and qPCR amplifications for all possible ligation sites using candidate primer pairs listed in Table S3.

### Chromatin Immunoprecipitation (ChIP) Assay

ChIP followed by allele-specific quantitative PCR (ChIP AS-qPCR) [38] were performed in U2OS cell line by using SimpleChIP^®^ Enzymatic Chromatin IP Kit (Magnetic Beads) (Catalog# 9003) (Cell Signaling Technology, USA) according to the manufacturer’s instruction. Briefly, approximately 30-50 million cells were cross-linked with 1% formaldehyde for 10 min and stopped the fixation reaction by quenching with 10 × glycine. DNA were digested with 0.4 μl of Micrococcal nuclease per tube for 20 min at 37°C with frequent mixing every 3 min, followed by stopping the digestion with 0.5M EDTA. The nuclear pellet was incubated in ChIP buffer with protease inhibitor cocktail for 10 min, followed by the sonication with 3 sets of 5 sec pulses and 30 sec pauses using an ultrasonic sonicator. The cross-linked chromatin were immunoprecipitated with ChIP grade Evi1 antibody (Santa Cruz Biotechnology, Texas, USA) or Histone H3 rabbit antibody as a positive control or normal rabbit immunoglobulin (*IgG*) as a negative control overnight at 4°C with rotation. Protein-DNA cross-links were precipitated by using ChIP-Grade Protein G Magnetic Beads. After reversal of protein-DNA cross-links by 5M NaCl and proteinase K, the DNA is purified using DNA purification spin columns. Following quantification was undertaken by ChIP AS-qPCR analysis with allele-specific primers listed in Table S3.

### DNA and RNA isolation and Quantitative Real-time PCR

The genomic DNA was isolated from the cells using Tiangen genomic DNA extraction Kit (Catalog#DP304, TianGen Biotech, China) according to the manufacturer’s instruction. The total RNA from the cells was isolated with TRIzol reagent (Invitrogen, USA), and 5 μg of total RNA per reaction were used to synthesize the complementary DNA (cDNA) with the Super Scripts II First-Strand cDNA synthesis kit (Invitrogen, USA). RT-qPCR was performed with QuantiTect SYBR Green PCR Kit (QIAGEN, Germany) according to the manufacturer’s instruction by CFX Connect^™^ Real-Time PCR Detection System (Bio-Rad, USA) with the primers listed in Table S3. Samples were tested in a 96-well format in triplicate. The housekeeping gene Glyceraldehyde 3-phosphate dehydrogenase (*GAPDH*) was used as an endogenous control to normalize the differences in treatment, amount of input cDNA and the amplification signal between the samples of different cell lines.

### Statistical Analysis

The mRNA expressions were calculated with 2^−ΔΔCt^ method as previously described [39]. Data were displayed as ± standard deviation (±SD) and unpaired, two-tailed student’s T test was used to calculate *P* value. Every analytical data were obtained from three independent experiments and each experiment was done in triplicate.

## Results

### Prioritizing rs13239597 as a potential functional independent variant through combining genomic fine-mapping and functional epigenetic analyses

To refine the independent association signals at 7q32.1 locus, we conducted a stepwise conditional association analysis using summarized GWAS data from 23,210 European samples [12]. Consistent with the previous study [8], we identified two independently associated signals within *IRF5* or *TNPO3* represented by 3 proxy SNPs (Figure 1). Adjusting the residual effect of 2 independent SNPs tagging association signal within *IRF5* retained 60 conditionally significant associated SNPs within *TNPO3* (*P* < 1 × 10^−5^, Figure 1). We further performed Bayesian fine-mapping analysis, resulting in 33 SNPs composing 95% credible set of this independent association signal (Figure 1). The majority (27/33) of credible SNPs are in strong LD (*r*^2^ > 0.8) with the most significant one rs12531054 (Figure 1). Particularly, we found that rs13239597 was the most significant SNP across the whole genome associated with both SLE and SSc from another large-scale pan-meta-analysis in 21,109 samples [8] (Figure S1), indicating its high potential functionality.

**Figure 1.**
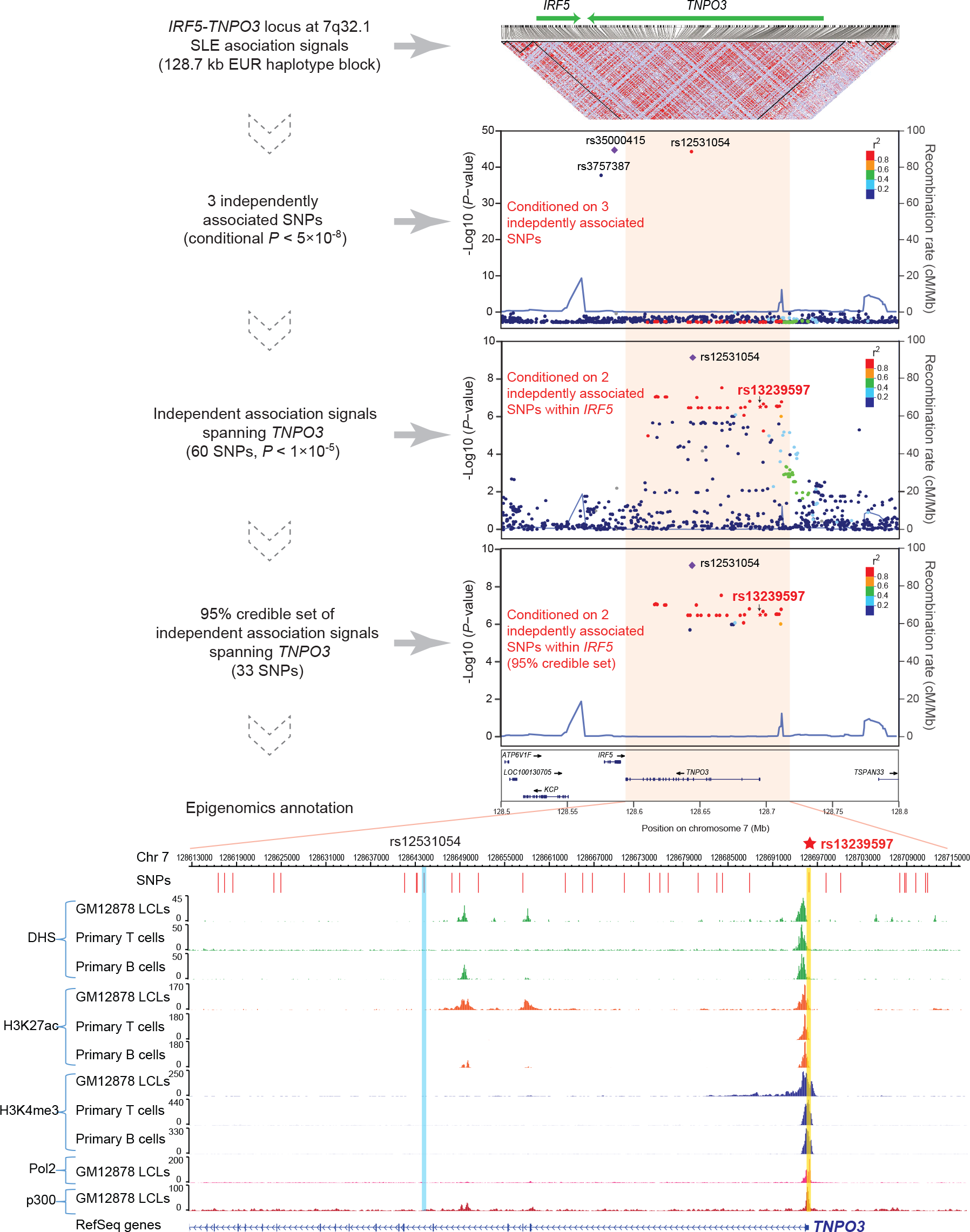
Prioritizing rs13239597 as a potential independent functional variant at 7q32.1 locus. The upper regional plots show the multi-step fine-mapping analysis results at 7q32.1 locus. The first inverted triangle shows haplotype block for all selected SNPs using 1000 genome-V3 European data [14]. The first regional plot shows stepwise conditional association analysis results, indicating 3 independently associated SNPs (p < 5 × 10^−8^) located at 7q32.1 locus. The middle regional plot shows conditional association signals after adjusting 2 of 3 independently associated SNPs near *IRF5*. The last regional plot shows Bayesian analysis [18] for the conditional association signals in the previous step. All regional plots are visualized using LocusZoom. The bottom part shows several active epigenetic annotation in three immunologically related blood cell lines for the independent associated region surrounding *TNPO3*. Related epigenetic data were visualized using WashU Epigenome Browser (v46.1), including DNase I hypersensitivity (DHS), the histone markers H3K27ac and H3K4me3, RNA polymerase II (Pol 2) and histone acetyltransferase p300 binding sites. The lead SNP rs12531054 region (pale blue) and our prioritized SNP rs13239597 region (pale yellow) are highlighted.

To evaluate the functionality of SNPs, we leveraged functional epigenetic annotation in several immunologically related blood cell lines. We found that rs13239597 was located near or within multiple extremely strong epigenetic enhancer markers in several immunologically related cell lines (GM12878 lymphoblastoid cells, primary T cells and B cells), including the DNase I hypersensitivity (DHS), the histone markers (H3K27ac, H3K4me3), RNA polymerase II (Pol 2) and histone acetyltransferase p300 binding sites (Figure 1), which strongly supported its potential functionality. By contrast, both the lead SNP rs12531054 as well as other credible SNPs were depleted of these functional epigenetic markers (Figure 1). Taken together, these analysis prioritized rs13239597 as a functional independent variant for further experimental validation.

### Identifying the candidate regulatory target gene of rs13239597

Rs13239597 was located in the promoter region of *TNPO3*. We performed cis-eQTL analysis using the data from 373 unrelated European samples on lymphoblastoid (LCLs) cell lines [14]. Among 21 surrounding genes detected, we found that the minor allele of rs13239597 (rs13239597-A) was exclusively and significantly associated with increased expression of *IRF5* (Bonferroni adjusted *P* = 1.742 × 10^−7^, Beta = 1.9) (Figure 2A and 2B; and Table 1). Consistent significantly reinforced effect of rs13239597-A allele on *IRF5* expression was observed on either LCLs from Genotype-Tissue Expression (GTEx) dataset [21] (*P* = 0.047, Beta = 0.28, Figure 2C and Table 1) or 5,311 peripheral blood samples from an eQTL meta-analysis [22] (*P* = 6.97 × 10^−21^, Z-score < −9.37, Table 1). By contrast, no significant association between rs13239597 and *TNPO3* was detected in either 373 LCLs or GTEx LCLs datasets (*P* > 0.05, Figure 2B and 2C; and Table 1). We further explored eQTL association between rs13239597 genotype and *IRF5* or *TNPO3* expression in other GTEx tissues [21], and detected significantly increased effect of rs13239597-A allele on *IRF5* expression in 26 different tissues (*P* < 0.05, Beta > 0), including some immunologically related tissues such as whole blood (*P* = 6.495 × 10^−4^, Beta = 0.191, Table S1) and thyroid (*P* = 9.283 × 10^−4^, Beta = 0.248, Table S1). We also detected significant eQTL association between rs13239597 genotype and *TNPO3* expression in 6 different GTEx tissues with discordant effect direction (*P* < 0.05, Beta > 0 for 3 tissues and Beta < 0 for another 3 tissues), including the whole blood (*P* = 2.481 × 10^−4^, Beta = −0.153, Table S1). Collectively, these eQTL results intensely highlighted *IRF5* as the candidate distal regulatory target gene of rs13239597.

**Table 1.** Comparison of Cis-eQTL analysis results of rs13239597 from three datasets.

| Gene | 373 LCLs<sup>a</sup> |  | GTEx LCLs<sup>b</sup> |  | Meta-analysis<sup>c</sup> |  |
| --- | --- | --- | --- | --- | --- | --- |
|  | *P* value | Beta | *P* value | Beta | *P* value | Z-score |
| *IRF5* | $1.742 \times 10^{-7}$ | 1.900 | 0.047 | 0.276 | $6.97 \times 10^{-21}$ | -9.37 |
| *TNPO3* | 0.714 | -0.111 | 0.589 | -0.076 | NA | NA |

Note: Cis-eQTL analysis results of rs13239597 for only *IRF5* and *TNPO3* expression were demonstrated in the table. <sup>a</sup> The data were obtained from 373 unrelated European samples on lymphoblastoid cell lines [14]. <sup>b</sup> The data were obtained from cells EBV-transformed lymphocytes of GTEx dataset [21]. <sup>c</sup> The data were obtained from peripheral blood samples of 5,311 individuals [22]. eQTL results for *TNPO3* expression in meta-analysis were not available (NA).

**Figure 2.**
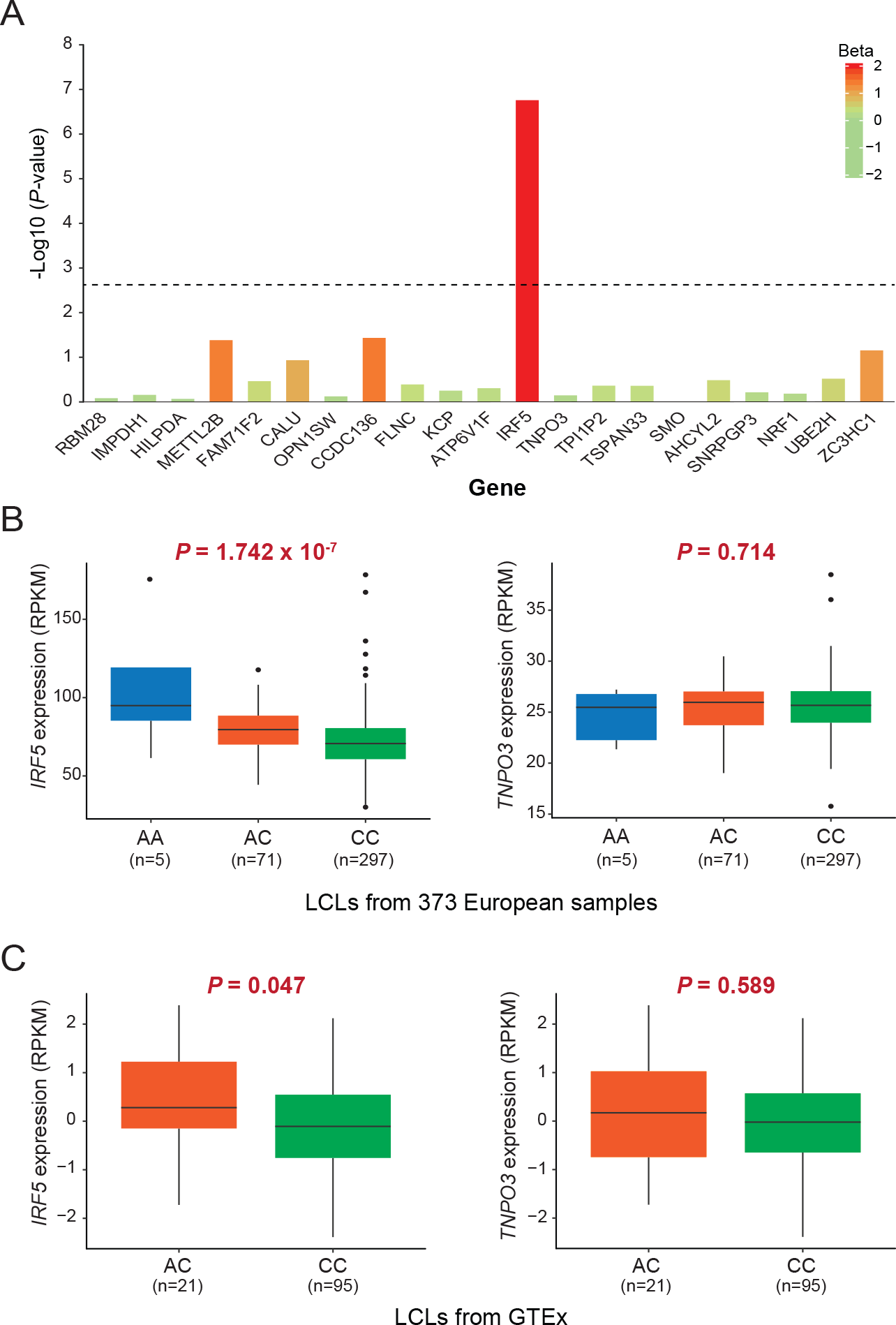
eQTL analysis demonstrates *IRF5* as a distal regulatory target gene for rs13239597. (A) Cis-eQTL analysis between rs13239597 genotype and 21 nearby genes (within 1000 kb) in lymphoblastoid (LCLs) cell lines from 373 unrelated samples [14]. The beta value represents the effect size for minor allele of rs13239597, and the dashed line represents the significant level after Bonferroni adjustment (Bonferroni adjusted *P* < 0.05). (B and C) Box plots show the comparison of *IRF5* or *TNPO3* expression with different genotypes (AA, AC and CC) of rs13239597 in lymphoblastoid (LCLs) cell lines from 373 unrelated European samples[14] (B) and in EBV-transformed lymphoblastoid cell lines (C) from GTEx[21]. The eQTL *P* values and sample count numbers (n) are shown.

### Validating the long-range chromatin interactions between rs13239597 and *IRF5*

The SNP rs13239597 was located ~118 kb far away from its candidate target gene *IRF5*. We therefore explored the potential long-range chromatin interactions between rs13239597 and *IRF5* using capture Hi-C and Hi-C data across multiple blood cell lines [23–26]. We observed that rs13239597 strongly interacted with the distal gene *IRF5* in five different cell lines (GM12878, CD34, K562, IMR90 and MCF7) (Figure 3A and Table S2). We also analyzed topologically associating domains (TADs) surrounding rs13239597 locus, and found that rs1323957-*IRF5* circuit was located within a conserved TAD with a size of 1.16 Mb in IMR90 cells (Figure 3B), further supported the distal chromatin interactions between them.

**Figure 3.**
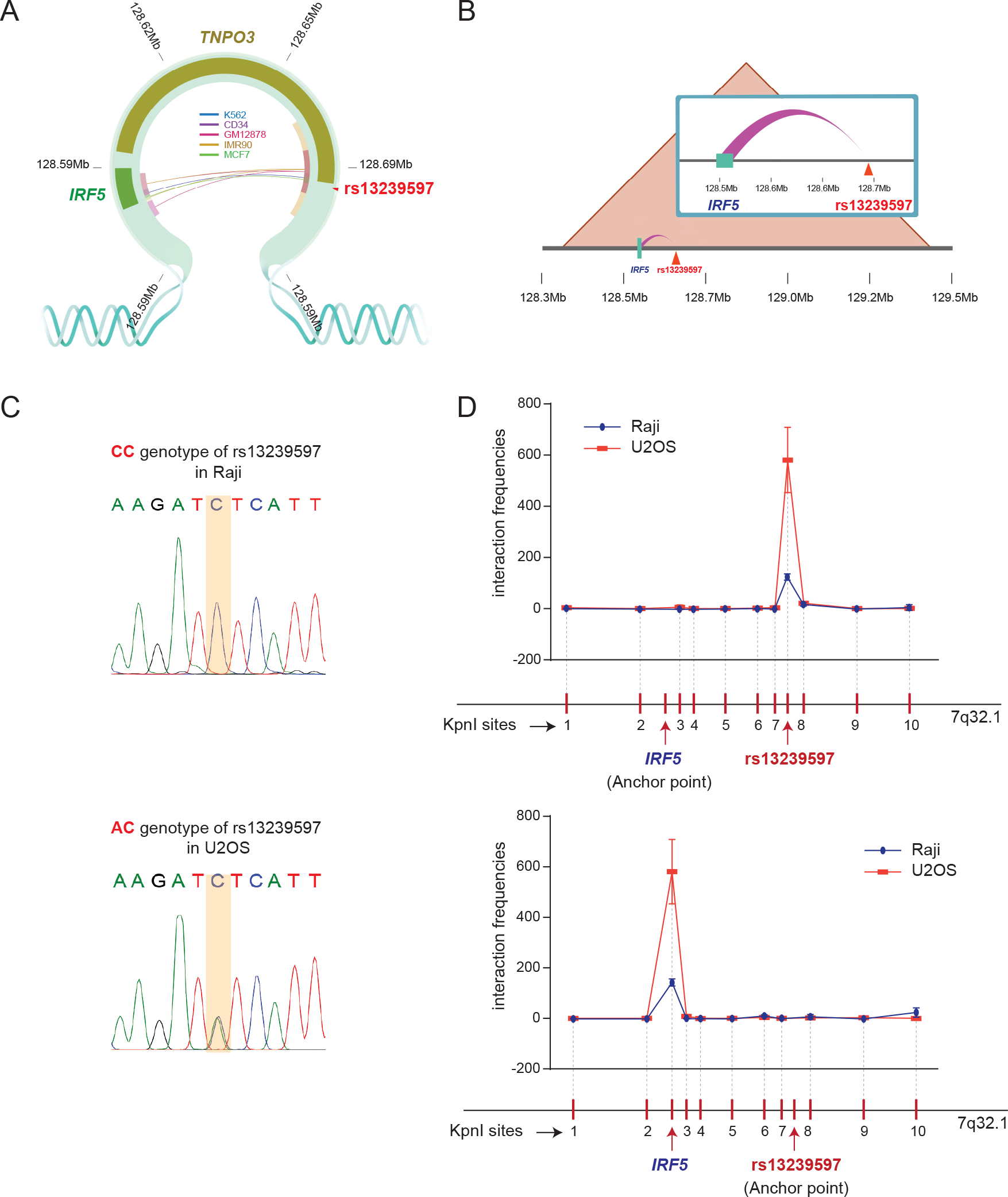
Validation of direct long-range chromatin interaction between rs13239597 and *IRF5*. (A) Hi-C interaction between rs13239597 and its distal target gene *IRF5*. Different colors of interacted lines between rs13239597 and *IRF5* indicate different five cell lines including K562, MCF7, IMR90, CD34 and GM12878 cell lines. (B) The looping interaction between rs13239597 and *IRF5* is located within the same topologically associated domain (TAD) with a size of 1.16 Mb in IMR90 cells. (C) Genotyping results of rs13239597 in Raji and U2OS cell lines. (D) Chromatin conformation capture (3C) assay in Raji cells (dark blue color line) and U2OS cells (orange color line). The interaction frequencies are shown between the region including *IRF5* promoter as the anchor point (upper) or the region harboring rs13239597 as the anchor point (lower) with other 10 neighboring KpnI sites (~100 kb upstream of *IRF5* and ~100 kb downstream of rs13239597) as the negative controls. Error bars are standard deviation (SD). Data are obtained from at least three replicates (n ≥ 3).

Genotyping assay revealed that B lymphocyte-derived Raji cell line is homozygous CC genotype for rs13239597 and U2OS cell line is heterozygous AC genotype for rs13239597 (Figure 3C). To directly validate the chromatin interactions between rs13239597 and *IRF5*, we performed 3C assay in Raji cell line. When anchored at *IRF5* promoter, rs13239597 enhancer region showed the strongest interaction with *IRF5* promoter region, compared with any of the other 10 neighboring KpnI sites tested (Figure 3D). Consistently, when anchored at rs13239597 enhancer region, *IRF5* promoter region showed the strongest interaction with rs13239597 enhancer region in comparison with other sites tested (Figure 3D). We also performed 3C assay in U2OS cell line, and detected relatively higher chromatin interactions between rs13239597 and *IRF5* compared with Raji cell line, which might indicate the superior allele-specific long-range chromatin loop formation between rs13239597-A and *IRF5*.

### Rs13239597 acts as an allele-specific enhancer regulating *IRF5* expression independent of *TNPO3*

To experimentally validate the allelic regulation between rs13239597 and *IRF5*, we compared the regulatory activity of genomic fragment containing different genotypes of rs13239597 on *IRF5* expression using dual-luciferase reporter assays in Raji cell line. We found that both alleles of rs13239597 could reinforce *IRF5* expression (*P* < 0.01, Figure 4A). Particularly, rs13239597-A allele had significantly enhanced effect on *IRF5* expression compared with rs13239597-C allele (Fold change = 2.06, *P* < 0.008, Figure 4A), which is consistent with our eQTL analysis results (Figure 2 and Table 1). Genotyping assay revealed that HEK293T cell line is homozygous CC genotype for rs13239597 (Figure S2). We further replicated luciferase assays in HEK293T cell lines. Consistent results were acquired in HEK293T cells in which significantly higher *IRF5* expression was observed on rs13239597-A allele compared with rs13239597-C allele (Fold change = 1.5, *P* < 0.01, Figure 4B). However, no significant enhanced luciferase activity was observed on rs13239597-C allele compared with the *IRF5* promoter-only plasmid (*P* > 0.05, Figure 4B). We also compared the luciferase activity of rs13239597-C or rs13239597-A allele on the nearby gene *TNPO3* in HEK293T cells, and detected no significant effect of rs13239597-A allele on *TNPO3* expression although different regulatory activities between two alleles of rs13239597 were detected (*P* < 0.02, Figure 4B). Collectively, these luciferase results suggested the allelic strong enhancer activity of rs13239597-A allele on *IRF5* instead of *TNPO3*.

**Figure 4.**
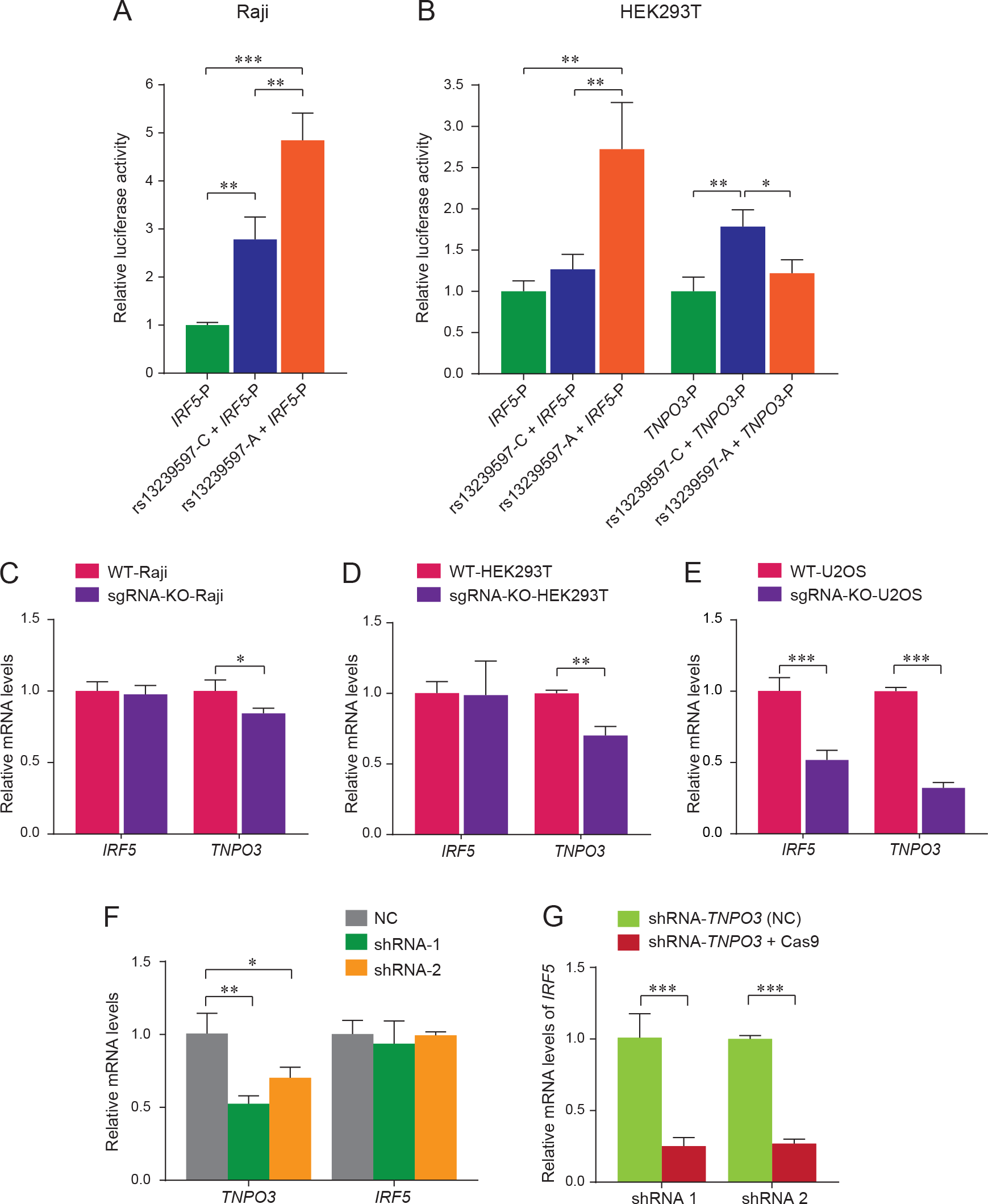
Validation of allele-specific enhancer activity of rs13239597 on *IRF5* expression. (A) Dual-luciferase reporter assays in Raji. The pGL3-basic vectors were constructed with *IRF5* promoter region and the region surrounding rs13239597-C or rs13239597-A, respectively. Luciferase signals are normalized to *Renilla* signals. (B) Dual-luciferase reporter assays in HEK293T cell lines. The pGL3-basic vectors were constructed with *IRF5* promoter region or *TNPO3* promoter region and the region surrounding rs13239597-C or rs13239597-A, respectively. Luciferase signals are normalized to *Renilla* signals. (C-E) Effect of deletion of the region residing rs13239597 by CRISPR-cas9 on *IRF5* and *TNPO3* expression in Raji (C), HEK293T (D) and U2OS (E) cell lines, respectively. Non-treated Raji, HEK293T and U2OS wild type (WT) cell lines are used as controls. (F) Effect of *TNPO3* knockdown on *IRF5* expression. Two independent shRNAs (shRNA-1 and shRNA-2) are used. shRNA-NC is used as the negative control. (G) Effect of deletion of the region residing rs13239597 by CRISPR-cas9 on *IRF5* on the basis of *TNPO3* knockdown in U2OS cell lines. Two independent shRNAs (shRNA-1 and shRNA-2) are used. Error bars, SD. n ≥ 3. \**P* ≤ 0.05, \*\**P* ≤ 0.01, \*\*\**P* ≤ 0.001 are determined by unpaired, two-tailed student’s T test. The samples are normalized by housekeeping gene *GAPDH*.

To further validate the allelic enhancer activity of rs13239597 on *IRF5*, we deleted the genomic fragment harboring rs13239597 using CRISPR-Cas9 in three different cell lines (Raji, HEK293T and U2OS). We observed no significant alterations of *IRF5* expression in the rs13239597-CC deleted Raji and HEK293T cells as compared with wild type cells (Figure 4C and 4D). By contrast, we detected significantly decreased *IRF5* expression in the rs13239597-AC deleted U2OS cell lines (*P* < 0.001, Figure 4E), supporting the allele-specific enhancer activity of rs13239597-A on *IRF5*.

We also detected significantly decreased expression of *TNPO3* in rs13239597-CC deleted cell lines (Figure 4C-4E), raising the question whether the detected regulatory activity of rs13239597 on *IRF5* was fictitious due to the intermediary effect of *TNPO3*. To further demonstrate the direct enhancer activity of rs13239597-A on *IRF5* independently of *TNPO3*, we firstly suppressed *TNPO3* expression in rs13239597-AC U2OS cells, and found no significant effect on *IRF5* expression (Figure 4F). We further deleted rs13239597-AC enhancer fragment in *TNPO3* suppressed U2OS cells, and detected significant decline of *IRF5* expression (*P* < 0.001, Figure 4G). Taken together, these experimental results prominently pinpointed that rs13239597 acted as allele-specific enhancer regulating *IRF5* expression independently of *TNPO3*.

### EVI1 preferentially binds to rs13239597-A to augment *IRF5* expression

We next explored the functional mechanism underlying rs13239597 as the strong allele-specific enhancer on *IRF5*. The motif analysis revealed that EVI1 could specifically bind to rs13239597-A allele (Figure 5A). To validate allelic binding affinity of EVI1 on rs13239597, we performed chromatin immunoprecipitation allele-specific quantitative PCR (ChIP AS-qPCR) assay in rs13239597-AC U2OS cell line. We found that EVI1 could bind to rs13239597 region in U2OS (Figure 5B). Particularly, EVI1 was preferentially recruited to the rs13239597-A allele compared with rs13239597-C allele (Fold change = 1.8, *P* < 0.001, Figure 5B). To further assess the transcriptional influence of rs13239597 on *IRF5* expression mediated by EVI1, we suppressed EVI1 by shRNA in both U2OS cells and HEK293T cells. Compared with shRNA NC transfected cell lines, we detected significant decline of *IRF5* expression in U2OS cells (*P* < 0.05, Figure 5C) while no obvious disturbance of *IRF5* expression in HEK293T cells (Figure 5D). We further co-transfected the EVI1 shRNA plasmids with rs13239597-A allele or rs13239597-C allele luciferase plasmids in HEK293T cells, and found that rs13239597-A allele significantly diminished *IRF5* expression (P < 0.01, Figure 5E) in contrast to that rs13239597-C allele had no decreased *IRF5* expression (Figure 5E), providing additional evidence for the allele-specific binding affinity of EVI1 to rs13239597-A. Taken together, we demonstrated that strong allele-specifically binding of EVI1 at the sequence harboring rs13239597-A allele could enhance expression of *IRF5*.

**Figure 5.**
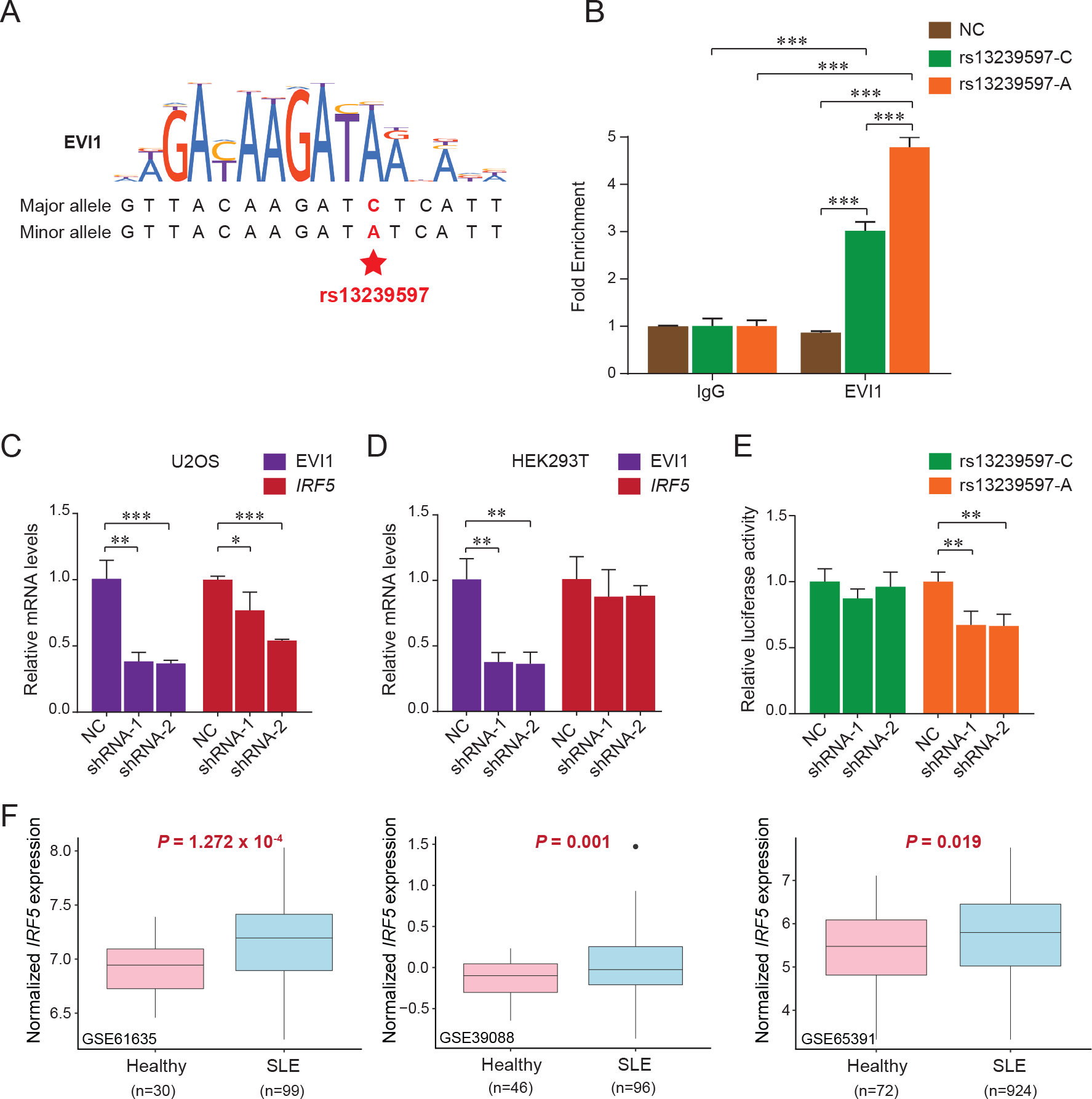
Preferential binding of EVI1 to rs13239597-A allele and increased *IRF5* expression in SLE patients. (A) EVI1 motif analysis for the sequences surrounding rs13239597. (B) Allele-specific ChIP assay for the comparison of EVI1 binding between rs13239597–C allele and rs13239597-A allele in U2OS cell line. Primers specifically targeting to rs13239597-C or rs13239597-A or *RPL30* exon (NC) region are used. The binding efficiency of EVI1 is shown as fold enrichment over *IgG*. (C and D) Effect of EVI1 knockdown on *IRF5* expression in U2OS (C) or HEK293T (D), respectively. Two independent shRNAs (shRNA-1 and shRNA-2) are used. The samples are normalized by housekeeping gene *GAPDH*. (E) Dual-luciferase reporter assay containing rs13239597-C allele or rs13239597-A allele plasmids co-transfected with two independent shRNA knockdown of EVI1 (shRNA-1 or shRNA-2). The shRNA-NC is used as the negative control. Luciferase signals are normalized to *Renilla* signals (n = 3). (F) Comparison of *IRF5* expression between healthy individuals and SLE patients in whole blood samples from three SLE genome-wide gene expression datasets (GSE61635, GSE39088 and GSE65391) [34] are shown. Error bars, SD. n ≥ 3. \**P* ≤ 0.05, \*\**P* ≤ 0.01, \*\*\**P* ≤ 0.001 are determined by unpaired, two-tailed student’s T test.

### Differential expression of *IRF5* between SLE patients and healthy individuals

To compare the *IRF5* expression between SLE patients and healthy individuals, we screened the *IRF5* expression in whole blood samples of three SLE genome-wide gene expression datasets (GSE61635, GSE39088 and GSE65391) [34]. Significantly higher *IRF5* expression was found in SLE patients compared with healthy individuals in all three datasets (*P* = 1.272 × 10^−4^ in GSE61635, *P* = 0.001 in GSE39088 and *P* = 0.019 in GSE65391, Figure 5F), which was consistent with that the risk allele-A of rs13239597 could augment *IRF5* expression.

## Discussion

GWASs have identified numerous SLE and SSc disease risk variants at *IRF5-TNPO3* locus of 7q32.1, however, the underlying functional mechanisms remain largely exclusive. Here, through the combination of a set of bioinformatics analyses and various experimental assays, we demonstrated that rs13239597 located in *TNPO3* promoter region functions as an allele-specific strong enhancer to directly modulate the distal gene *IRF5* by long-range chromatin loop formation. We corroborated that the enhancer activity of rs13239597 was elevated by the transcription factor EVI1 preferentially recruited to rs13239597-A allele, Which efficiently enhanced the *IRF5* expression. Taken together, our study highlighted the allele-specific functional role of rs13239597 leading to the *IRF5* hyperactivation followed by the production of pathogenic antibodies against self-tissues (Figure 6A).

**Figure 6.**
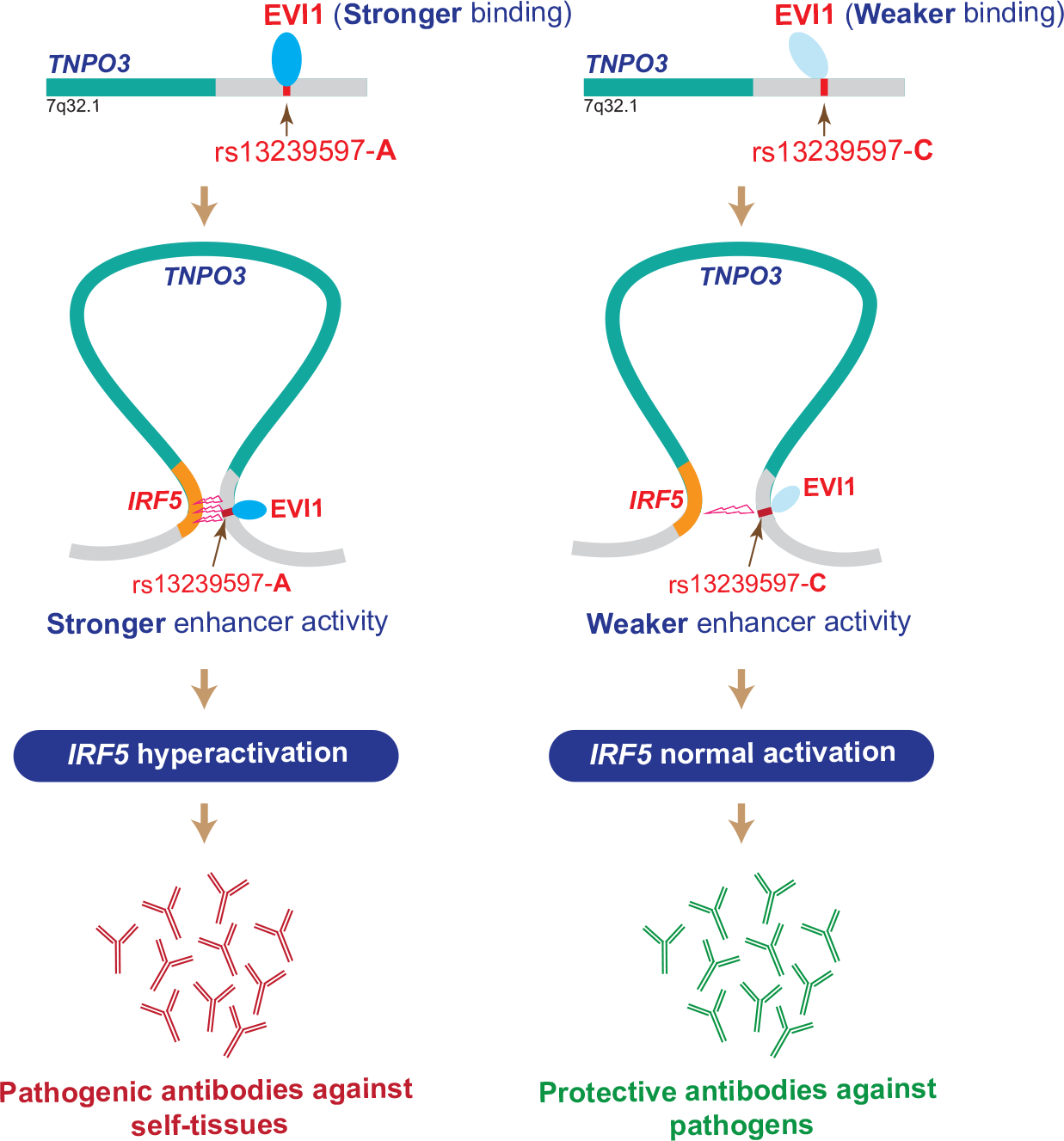
A schematic proposed model between rs13239597 and *IRF5*. The schematic model shows how a noncoding variant (rs13239597) influences the autoantibody production. The left panel shows rs13239597-A allele vigorously binds to EVI, which leads to *IRF5* hyperactivation via long-range chromatin loop formation, resulting in pathogenic autoantibody production against self-tissues. In contrast, the right panel shows rs13239597-C allele has a weaker activity of binding to EVI, which leads to *IRF5* normal activation, resulting in the protective autoantibody production against pathogens.

Our analysis showed that a GWAS risk variant associated with both SLE and SSc could regulate *IRF5* expression via long-range loop formation. *IRF5* is a member of interferon regulatory factor (IRF) family which is the transcriptional regulators of Type I interferons (IFNs) playing crucial role in modulating inflammatory immune responses in numerous cell types, including dendritic cells, macrophages and B cells [40]. Studies on murine models also indicated that *Irf*^−/−^ mice could result in poor lymphocyte activation, decreased autoantibody levels [41] and significant lower production of IFN-I and key cytokines (IL-12 and IL-23) that link innate immunity to SLE pathogenesis [42]. Consistently, our analysis demonstrated that the risk allele of rs13239597 could augment *IRF5* expression, and *IRF5* was significantly higher expressed in SLE patients compared with healthy individuals [34].

Regulation of mammalian gene transcription is accomplished by proximal promoter and distal enhancer, which had noticeable functional similarities such as DNase I hypersensitivity, histone modification patterns and transcription factor binding sites [43]. Dao *et al.* recently verified the functional mechanisms of Epromoters referred to as the promoters with enhancer functions, suggesting that regulatory elements with dual roles as transcriptional promoters and enhancers were strongly involved in *cis* regulation of distal gene expression in natural context [44]. Recently, the functional SNPs with both promoter and enhancer activities were reported [45], in which different alleles of a SNP mechanistically perform different functions. In our study, rs13239597 located in the promoter region of *TNPO3* functions as an allele-specific functional enhancer, which highlights dual roles of its functionality.

Our study revealed that the transcription factor EVI1 could preferentially bind to rs13239597-A allele to increase *IRF5* expression. EVI1 is crucial for hematopoietic stem cells originating in bone marrow [46], which give rise to production of human lymphoid cells including T cells, B cells and natural killer cells [47]. Many IRF family members play important roles in the differentiation of hematopoietic cells [48]. A recent SLE and SSc combined meta-analysis revealed that the minor rs13239597-A allele could increase disease risk (OR = 1.848 for SLE and OR = 1.567 for SSc) [8]. Consistently, our study revealed that EVI1 strongly binds to the risk allele (rs13239597-A) and subsequently promotes *IRF5* expression, high expression of which produces pathogenic antibodies leading to SLE and SSc.

In summary, we provided a new mechanistic insight that rs13239597 acts as an allele-specific strong enhancer to directly regulate *IRF5* expression mediated by EVI1. We established the feasible approach to investigate the role of a noncoding functional variant and its target distal gene through a series of integrated bioinformatics data analyses and various functional assays. We anticipate that the similar approach could be the further investigation of functional mechanisms underlying disease risk variants associated with more human complex diseases to fulfill the gap of GWASs. We believe that our findings might fulfill the current issues towards the understanding of complex genetic architecture and the promising therapeutic target for both SLE and SSc autoimmune diseases.

## Supplemental Data

Supplemental Data include two figures and three tables.

## Web Resources

1000 Genomes V3 genotype data, ftp://ftp.trace.ncbi.nih.gov/1000genomes/ftp/release/20130502/

4DGenome, https://4dgenome.research.chop.edu/

ArrayExpress (E-GEUV-1), http://www.ebi.ac.uk/arrayexpress/experiments/E-GEUV-1/

CRISPR design platform (V1 tool), http://crispr.mit.edu/

dbGaP, https://www.ncbi.nlm.nih.gov/gap

GEO, https://www.ncbi.nlm.nih.gov/gds/

GTEx Portal, https://www.gtexportal.org/home/

GWAS Catalog, http://www.ebi.ac.uk/gwas/

HaploReg, http://www.broadinstitute.org/mammals/haploreg/haploreg.php

LocusZoom, http://locuszoom.sph.umich.edu/

MEME Suite, http://meme-suite.org/

OMIM, http://www.omim.org/

PLINK, http://zzz.bwh.harvard.edu/plink/

R statistical software, https://www.r-project.org/

SLE GWAS data, http://insidegen.com/insidegen-LUPUS-data.html

UCSC ENCODE download portal, https://genome.ucsc.edu/encode/downloads.html

Vcftools, http://vcftools.sourceforge.net/

WashU Epigenome Browser (v46.1), http://epigenomegateway.wustl.edu/browser/

## Acknowledgements

Not Applicable.

## Authors’ contributions

TLY and YG designed and supervised the project. HNT designed and performed the experiments and wrote the manuscript. XFC conducted the data analysis and revised the manuscript. WXH, YYD, DLZ, HC, NNW and HHC contributed to design the experiments. Other authors contributed to the manuscript preparation. All authors read and approved the final manuscript.

