## Supplementary materials for "An allele-specific functional SNP associated with two autoimmune diseases modulates *IRF5* expression by long-range chromatin loop formation"

**Supplemental Figures**

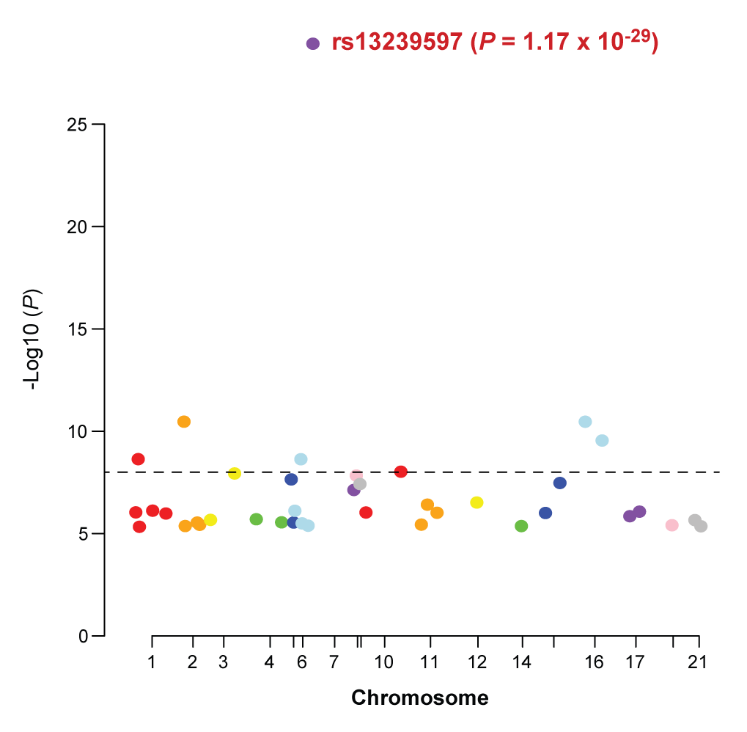

**Supplementary Fig. 1. rs13239597 is the most significantly associated SNP with SLE and SSc.**

Manhattan Plot shows the SNPs associated with SLE and SSc in the whole genome region (*P* < 1 × 10^-5^) downloaded the data from a pan-meta-analysis of two GWASs including SLE and SSc cohorts ^1^. The dashed line represents the significant level (*P* = 1 × 10^-8^).

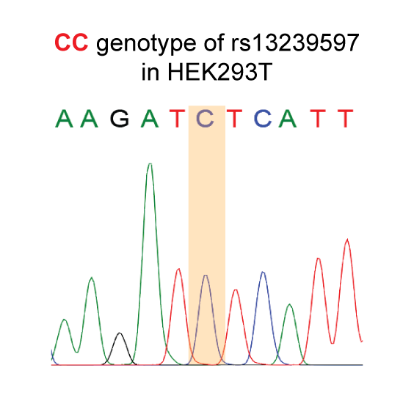

**Supplementary Fig. 2. Genotyping result of rs13239597 in HEK293T cell line.**

The color highlighted peak shows the homozygous CC genotype of rs13239597 in HEK293T cell line.

**Supplementary Tables**

**Supplementary Table 1. Cis-eQTL analysis results of rs13239597 for *IRF5* and *TNPO3* expression from Genotype-Tissue Expression (GTEx) dataset**

| GTEx Tissues | *IRF5* | | *TNPO3* | |
| --- | --- | --- | --- | --- |
|  | *P*-value | Beta | *P*-value | Beta |
| Adipose_Subcutaneous | **0.006** | **0.105** | 0.187 | -0.068 |
| Adipose_Visceral_Omentum | 0.165 | 0.079 | 0.547 | -0.033 |
| Adrenal_Gland | **0.016** | **0.213** | **0.044** | **-0.114** |
| Artery_Aorta | **0.025** | **0.124** | 0.095 | -0.089 |
| Artery_Coronary | **0.025** | **0.154** | 0.181 | -0.088 |
| Artery_Tibial | **8.581 × 10^-4^** | **0.178** | 0.195 | -0.050 |
| Brain_Amygdala | 0.275 | 0.153 | **0.004** | **0.443** |
| Brain_Anterior_cingulate_cortex_BA24 | **0.036** | **0.325** | 0.066 | 0.162 |
| Brain_Caudate_basal_ganglia | 0.359 | 0.105 | 0.333 | 0.104 |
| Brain_Cerebellar_Hemisphere | 0.065 | -0.304 | 0.840 | -0.018 |
| Brain_Cerebellum | **0.021** | **-0.355** | 0.430 | -0.066 |
| Brain_Cortex | 0.583 | 0.092 | 0.832 | 0.017 |
| Brain_Frontal_Cortex_BA9 | **0.028** | **0.380** | 0.288 | 0.084 |
| Brain_Hippocampus | 0.607 | 0.053 | 0.311 | 0.101 |
| Brain_Hypothalamus | 0.727 | -0.051 | 0.235 | 0.096 |
| Brain_Nucleus_accumbens_basal_ganglia | 0.816 | 0.033 | 0.476 | 0.066 |
| Brain_Putamen_basal_ganglia | 0.740 | 0.061 | 0.500 | 0.097 |
| Brain_Spinal_cord_cervical_c-1 | 0.486 | 0.114 | 0.118 | -0.352 |
| Brain_Substantia_nigra | 0.566 | 0.079 | 0.528 | 0.091 |
| Breast_Mammary_Tissue | 0.114 | 0.120 | 0.414 | 0.054 |
| Cells_EBV-transformed_lymphocytes | **0.047** | **0.276** | 0.589 | -0.076 |
| Cells_Transformed_fibroblasts | **0.045** | **0.236** | 0.451 | -0.030 |
| Colon_Sigmoid | 0.124 | 0.171 | 0.930 | 0.005 |
| Colon_Transverse | **0.007** | **0.231** | **0.003** | **0.153** |
| Esophagus_Gastroesophageal_Junction | **0.050** | **0.153** | 0.908 | 0.006 |
| Esophagus_Mucosa | **0.004** | **0.180** | 0.763 | 0.009 |
| Esophagus_Muscularis | **0.001** | **0.206** | 0.757 | 0.013 |
| Heart_Atrial_Appendage | **0.013** | **0.156** | 0.246 | 0.054 |
| Heart_Left_Ventricle | 0.103 | 0.135 | 0.412 | 0.031 |
| Liver | **0.032** | **0.297** | 0.446 | 0.060 |
| Lung | **0.003** | **0.135** | 0.919 | -0.005 |
| Minor_Salivary_Gland | 0.996 | -0.001 | 0.649 | 0.046 |
| Muscle_Skeletal | **1.957 × 10^-4^** | **0.245** | **0.027** | **-0.084** |
| Nerve_Tibial | **2.463 × 10^-4^** | **0.208** | 0.541 | 0.037 |
| Ovary | 0.669 | 0.053 | 0.534 | -0.092 |
| Pancreas | **0.004** | **0.284** | 0.233 | -0.067 |
| Pituitary | **0.011** | **0.363** | 0.611 | 0.046 |
| Prostate | **0.046** | **0.259** | 0.822 | 0.018 |
| Skin_Not_Sun_Exposed_Suprapubic | **0.027** | **0.156** | 0.968 | -0.002 |
| Skin_Sun_Exposed_Lower_leg | **4.796 × 10^-4^** | **0.207** | 0.341 | 0.041 |
| Small_Intestine_Terminal_Ileum | 0.093 | 0.128 | 0.454 | 0.067 |
| Spleen | 0.820 | 0.028 | 0.494 | -0.051 |
| Stomach | **0.025** | **0.208** | 0.446 | -0.050 |
| Thyroid | **9.283 × 10^-4^** | **0.248** | 0.606 | 0.028 |
| Uterus | 0.220 | -0.181 | **0.018** | **0.300** |
| Vagina | 0.170 | 0.175 | 0.127 | 0.147 |
| Whole_Blood | **6.495 × 10^-4^** | **0.191** | **2.481 × 10^-4^** | **-0.153** |

Note: The analyzed data were obtained from GTEx dataset ^2^. The significantly correlated tissues for *IRF5* and *TNPO3* expression (*P* < 0.05) were highlighted as bold pattern.

**Supplementary Table 2. Chromatin interactions between rs13239597 (Locus 1) and *IRF5* (Locus 2)**

| Dataset^a^ | Cell^b^ | Locus 1 | Locus 2 | References |
| --- | --- | --- | --- | --- |
| Capture Hi-C | GM12878 | chr7:128683895-128700441 | chr7:128579427-128588145 | Mifsud, B. et al. ^3^ |
| Capture Hi-C | GM12878 | chr7:128683895-128700441 | chr7:128570699-128576889 | Mifsud, B. et al. ^3^ |
| Capture Hi-C | CD34 | chr7:128683895-128700441 | chr7:128570699-128576889 | Mifsud, B. et al. ^3^ |
| Capture Hi-C | CD34 | chr7:128683895-128700441 | chr7:128579427-128588145 | Mifsud, B. et al. ^3^ |
| Hi-C (4D Genome) | IMR90 | chr7:128672324-128710090 | chr7:128579429-128588147 | Jin, F. et al. ^4^ |
| Hi-C (4D Genome) | IMR90 | chr7:128672694-128710090 | chr7:128576892-128579428 | Jin, F. et al. ^4^ |
| ChIA-PET^ENCODE^ | MCF7 | chr7:128694260-128696368 | chr7:128577672-128579301 | Harrow, J. et al. ^5^ |
| ChIA-PET^ENCODE^ | K562 | chr7:128692602-128696425 | chr7:128578528-128581641 | Harrow, J. et al. ^5^ |

Note: ^a^Dataset, Capture Hi-C, Hi-C or ChIA-PET data used; ^b^Cell, Hi-C data on human healthy cells or ChIA-PET data on all cells were collected; ChIA-PET^ENCODE^, ChIA-PET data retrieved from UCSC ENCODE download portal; Locus 1 and Locus 2, chromatin interactions regions (hg19).

**Supplementary Table 3. Primers used in different experiments**

| Experiments | Target name | Primers | |
| --- | --- | --- | --- |
|  |  | **Forward** | **Reverse** |
| Luciferase plasmid construction | Enhancer (rs13239597) | CGACGCGTATTCAACTCTCCCTAGAGTGGCCA (MluI) | TCCCCCGGGCTTCCCTTTCATCTAGCTTCTCCC (SmaI) |
|  | Promoter (*IRF5*) | TCCCCCGGGGTGTGGGTTTGCAAGGAGACATGT (SmaI) | CCCAAGCTTCTCTGAGCTGCTACTGTCTCACAT (HindIII) |
|  | Enh+Pro (*TNPO3*) | CGACGCGTGCTTCCCTTTCATCTAGCTTCTCCC (MluI) | CCCAAGCTTTGGTACACGAGCTGCAATGTC (HindIII) |
|  | Promoter (*TNPO3*) | CCGCTCGAGAGTGCAAAGGACTAGCAGCA (XhoI) | CCCAAGCTTTGGTACACGAGCTGCAATGTC (HindIII) |
|  | Site-directed mutagenesis | GATATCATTTAATCCGCCTAACAACCTTGCCA | GCGGATTAAATGATATCTTGTAACTACAGTGTTTTACACATTGTG |
| CRISPR-Cas9 | Upstream_sgRNA | ACCGCCTAGAGTGGCCACAGTAC | AAACGTACTGTGGCCACTCTAGG |
|  | Downstream_sgRNA | ACCGAGAAAAGTGGCCCATGCTC | AAACGAGCATGGGCCACTTTTCT |
|  | Nest-PCR_upstream | AAGGCTCTCCGTGGAAGTTG | AGTGCAAAGGACTAGCAGCA |
|  | Inner-PCR_upstream | GTACACTACAAACGGCCTCCT | AGCTGTAAGCAGAATGAAGACCT |
|  | Nest-PCR_downstream | AACCCGTTTTAGGCGCTGA | GGCCACAATAGGGTGCAGAT |
|  | Inner-PCR_downstream | AAGGTGATTTGTGCTCTCGC | TTCGGGCTTGAGGGAAACAC |
| shRNA knockdown | *TNPO3*-shRNA-1 | TGCTGTTGACAGTGAGCGACGGCGCACAGAAATTATAGAATAGTGAAGCCACAGATGTA | TCCGAGGCAGTAGGCACCGGCGCACAGAAATTATAGAATACATCTGTGGCTTCACTA |
|  | *TNPO3*-shRNA-2 | TGCTGTTGACAGTGAGCGACCTCAATATGAGGTAGTAGAATAGTGAAGCCACAGATGTA | TCCGAGGCAGTAGGCACCCTCAATATGAGGTAGTAGAATACATCTGTGGCTTCACTA |
|  | EVI1-shRNA-1 | TGCTGTTGACAGTGAGCGAGCACTACGTCTTCCTTAAATATAGTGAAGCCACAGATGTA | TCCGAGGCAGTAGGCACGCACTACGTCTTCCTTAAATATACATCTGTGGCTTCACTA |
|  | EVI1-shRNA-2 | TGCTGTTGACAGTGAGCGACCTTTCTTTATGGACCCTATTTAGTGAAGCCACAGATGTA | TCCGAGGCAGTAGGCACCCTTTCTTTATGGACCCTATTTACATCTGTGGCTTCACTA |
|  | shRNA-NC | TGCTGTTGACAGTGAGCGAGTTCTCCGAACGTGTCACGTTAGTGAAGCCACAGATGTA | TCCGAGGCAGTAGGCACGTTCTCCGAACGTGTCACGTTACATCTGTGGCTTCACTA |
|  | Mir-30 | CAGAAGGCTCGAGAAGGTATATTGCTGTTGACAGTGAGCG | GTAGCCCCTTGAATTCGTCGACTCCGAGGCAGTAGGCA |
| ChIP | A-allele-specific | AACATTGCTGCTAGTCCTTTGC | GTTAGGCGGATTAAATGAT |
|  | C-allele-specific | AACATTGCTGCTAGTCCTTTGC | GTTAGGCGGATTAAATGAG |
| 3C | Target site | CTCGCCCTGCTGTGTTCA | GTGGCACATATTTCAGGTTTGT |
|  | NC 1 | TCTTGTGGCAAAACTGCACC | CTCTTGTCATCCTCTACCCCAA |
|  | NC 2 | GCTGTAGGCACCCACCGAAGC | CAGTAGTCACCGGTAAGTGCC |
|  | NC 3 | GTGCAGAATGAACCACTTCCCTG | CTGTTCTGTCTGGTGTTGTAGCC |
|  | NC 4 | TGACTATCTTTAGGCATGGTGAAC | CCCCGTCAGGAGATCTGTCTA |
|  | NC 5 | GGCAGTGTCGAGCCAAGGCG | CGTGGGAAGACAATTTGGGGAG |
|  | NC 6 | CTCTGGCGGGCAGAAGTAAGGC | CCTTCATCAAGGAAGCAAGTGGC |
|  | NC 7 | GCTACGCACATGGATGAACCCTG | GTGCAACCATCACCATCATCC |
|  | NC 8 | CCCCACAGTCTGCTGAGTCCC | GGGCCTGCTGGATGCTCACAGG |
|  | NC 9 | TGCCAGGTGGAAGGGTAGAT | TGGGTAGCACTGAGTGTTTG |
|  | NC 10 | TTCCCTGGGACTTGGGAATTG | CTCCACTCCTTCCTTCTCTTGT |
| qRT-PCR | *IRF5* | GGGCTTCAATGGGTCAACG | GCCTTCGGTGTATTTCCCTG |
|  | *TNPO3* | AGCTAATCGGCGCACAGAAA | ACCAACTTCCCAAACAGCGA |
|  | EVI1 | CATTGGGAACAGCAACCAT | GGTCACCAAAGCCTTTTCAT |
|  | *GAPDH* | GGAGCGAGATCCCTCCAAAAT | GGCTGTTGTCATACTTCTCATGG |
